# The Dreem Headband as an Alternative to Polysomnography for EEG Signal Acquisition and Sleep Staging

**DOI:** 10.1101/662734

**Authors:** Pierrick J. Arnal, Valentin Thorey, Michael E. Ballard, Albert Bou Hernandez, Antoine Guillot, Hugo Jourde, Mason Harris, Mathias Guillard, Pascal Van Beers, Mounir Chennaoui, Fabien Sauvet

## Abstract

Despite the central role of sleep in our lives and the high prevalence of sleep disorders, sleep is still poorly understood. The development of ambulatory technologies capable of monitoring brain activity during sleep longitudinally is critical to advancing sleep science and facilitating the diagnosis of sleep disorders. We introduced the Dreem headband (DH) as an affordable, comfortable, and user-friendly alternative to polysomnography (PSG). The purpose of this study was to assess the signal acquisition of the DH and the performance of its embedded automatic sleep staging algorithms compared to the gold-standard clinical PSG scored by 5 sleep experts. Thirty-one subjects completed an over-night sleep study at a sleep center while wearing both a PSG and the DH simultaneously. We assessed 1) the EEG signal quality between the DH and the PSG, 2) the heart rate, breathing frequency, and respiration rate variability (RRV) agreement between the DH and the PSG, and 3) the performance of the DH’s automatic sleep staging according to AASM guidelines vs. PSG sleep experts manual scoring. Results demonstrate a strong correlation between the EEG signals acquired by the DH and those from the PSG, and the signals acquired by the DH enable monitoring of alpha (r= 0.71 ± 0.13), beta (r= 0.71 ± 0.18), delta (r = 0.76 ± 0.14), and theta (r = 0.61 ± 0.12) frequencies during sleep. The mean absolute error for heart rate, breathing frequency and RRV was 1.2 ± 0.5 bpm, 0.3 ± 0.2 cpm and 3.2 ± 0.6 %, respectively. Automatic Sleep Staging reached an overall accuracy of 83.5 ± 6.4% (F1 score : 83.8 ± 6.3) for the DH to be compared with an average of 86.4 ± 8.0% (F1 score: 86.3 ± 7.4) for the five sleep experts. These results demonstrate the capacity of the DH to both precisely monitor sleep-related physiological signals and process them accurately into sleep stages. This device paves the way for high-quality, large-scale, longitudinal sleep studies.

## Introduction

Sleep disorders and insufficient sleep negatively impact hundreds of millions of people across the world and constitute a growing public health epidemic with grave consequences, including increased risk of cardiovascular and neurodegenerative diseases and psychiatric disorders (1). The most prevalent sleep disorders include insomnia, which affects ~20% of the general population, and obstructive sleep apnea, which affects ~10% of the general population (2). Despite their high prevalence, sleep disorders remain largely unidentified and/or untreated with less than 20% of patients estimated to be accurately diagnosed and treated (3).

Today, the gold standard to study or diagnose sleep disorders is nocturnal polysomnography (PSG). A PSG sleep study is typically a single overnight assessment, usually taking place in a sleep center, during which physiological signals including electroencephalographic (EEG), electromyographic (EMG), and electrooculographic (EOG) activity, breathing effort, airflow, pulse, and blood oxygen saturation are recorded. Analysis of these signals relies on trained sleep experts to visually inspect and manually annotate and recognize specific EEG, EOG, EMG patterns on 30-second segments (epochs) of the full PSG recording to score sleep stages (Wake, sleep stages 1 (N1), 2 (N2) and 3 (N3), and REM sleep), according to the American Academy of Sleep Medicine’s [AASM] guidelines (4)).

However, the gold-standard PSG suffers from several limitations. From a practical standpoint, a PSG is complicated and time-consuming to set-up, requiring up to one hour to install by a trained sleep technician; it is also quite expensive (typically $1,500-$2,000 per night in the US). Furthermore, a clinical PSG may not reliably capture a patient’s typical sleep because it is cumbersome and the clinical setting often generates stress for the patient. Moreover, because a PSG is generally performed over only one night, it does not capture intra-individual variability across nights and the final diagnosis is often rendered on an unrepresentative night of sleep (5, 6). From a clinical standpoint, the way PSG records are analyzed by sleep experts is unsatisfactory: it requires extensive training, it is time-consuming, inconsistent, and suffers from poor inter-rater reliability. For instance, one study conducted on the AASM ISR data set found that sleep stage agreement across experts averaged 82.6% using data from more than 2,500 scorers, most with 3 or more years of experience, who scored 9 record fragments, representing 1,800 epochs (i.e. more than 3,200,000 scoring decisions). The agreement was highest for the REM sleep stage (90.5%) and slightly lower for N2 and Wake (85.2% and 84.1%, respectively), while the agreement was far lower for stages N3 and N1 (67.4% and 63.0%, respectively), placing constraints on the reliability of manual scoring (7). Critically, studies also indicate that agreement varies substantially across different sleep pathologies and sleep centers (7, 8).

Automatic PSG analysis in sleep medicine has been explored and debated for some time, but has yet to be widely adopted in clinical practice. In recent years, dozens of algorithms have been published that achieve expert-level performance for automated analysis of PSG data (9–12). Indeed, scientists and engineers have used artificial intelligence (AI) methods to develop automated sleep stage classifiers and EEG pattern detectors, thanks to open access sleep data sets such as the National Sleep Research Resource (https://sleepdata.org). Regarding sleep staging, Biswal et al. proposed the SLEEP-NET algorithm (13), a deep recurrent neural network trained on 10,000 PSG recordings from the Massachusetts General Hospital Sleep Laboratory. The algorithm achieved an overall accuracy comparable to human-level performance of 85.76% (N1 : 56%, N2 : 88%, N3 : 85%, REM : 92%, Wake : 85%). Another important collaborative study recently published an algorithm validated on ~3,000 normal and abnormal sleep recordings. They showed that their best model using a deep neural network performed better than any individual scorer (overall accuracy: 87% compared to the consensus of 6 scorers). The problem of low inter-scorer reliability of sleep stages is addressed by using a consensus of multiple trained sleep scorers instead of relying on a single expert’s interpretation (8, 12, 14). Regarding the topic of sleep EEG event detection, deep learning methods have shown state-of-the-art performance for automatic detection of sleep events such as spindles and k-complexes in PSG records (15).

With the rise of wearable technology over the last decade, consumer sleep trackers have seen exponential growth (16). For many years, these devices used only movement analysis, called actigraphy, before incorporating measures of pulse oximetry. Actigraphy has been extensively used in sleep research for sleep-wake cycle assessment at home. However, this measure has very low specificity for differentiating sleep from motionless wakefulness, resulting in an overestimation of total sleep time (TST) and underestimation of wake after sleep onset (WASO) time (17, 18). Thus, actigraphy is still quite far from being a reliable alternative to PSG. And though the addition of pulse oximetry improves analysis over actigraphy alone, it still only enables rough estimations of sleep efficiency and stages. This is because the essential component of monitoring brain electrical activity with EEG sensors was still lacking.

More recently, a new group of devices has emerged for home sleep monitoring that uses EEG electrodes to measure brain activity. These include headbands (19–22) and devices placed around the ear (23, 24). Unlike traditional PSG, these more compact devices are usually cheaper, less burdensome, designed to be worn for multiple nights at home to enable longitudinal data collection, and require minimal or no expert supervision. Only a few of these device makers have published their performance compared to PSG; and those that have often only report aggregated metrics rather than raw data, and do not permit open access to the data set so that results can be independently verified. Perhaps most notably, however, this new generation of home sleep trackers generally suffers from only mediocre accuracy and reliability compared to the gold standard PSG.

In this study, we introduce the Dreem headband (DH) which is intended as an affordable, comfortable, and user-friendly alternative to PSG with a high level of accuracy regarding both physiological signal acquisition and automatic sleep stage analysis using a deep learning algorithm along with 5 dry EEG electrodes (O1, O2, FpZ, F7, F8), a 3-D accelerometer, and a pulse oximeter embedded in the device. To this end, we recorded data from 25 subjects over a single night using the DH and a clinical PSG simultaneously. We assessed: 1) the EEG signal quality and the ability of the DH to monitor brain sleep frequencies during the night; 2) the accuracy of heart rate, breathing frequency, and respiration rate variability (RRV) during sleep and 3) the performance of the automatic sleep stage classification algorithm of the DH compared to a consensus of 5 sleep experts’ manual scoring of the PSG. The data set of the current study is available from the corresponding author on reasonable request.

## Methods

### Subjects

Thirty-one volunteers were recruited without regard to gender or ethnicity from the local community by study advertisement flyers. Volunteers were eligible if they were between the ages of 18 and 65 years and capable of providing informed consent. Exclusion criteria included current pregnancy or nursing; severe cardiac, neurological, or psychiatric comorbidity in the last 12 months; morbid obesity (BMI >= 40); or use of benzodiazepines, non-benzodiazepines (Z-drugs), or gammahydroxybutyrate (GHB) on the day of the study.

Each participant provided one night of data; with the exception of 2 participants who completed a second night each due to data loss related to PSG battery issues on their first nights of study. In total, 8 nights of data were excluded from the final analysis data set: 3 due to poor signal or system malfunction including battery issues on PSG, 2 due to the discovery of asymptomatic Apnea–Hypopnea Index (AHI) > 5 during the course of the study, and 1 due to an unusually short total sleep time (4.5 hours).

The final analysis data set consisted of one night record from each of 25 participants; demographics are summarized in Table 1. The sample included individuals with self-reported sleep quality ranging from no complaints to sub-threshold insomnia symptoms and moderate to severe daytime sleepiness. Only one met the Insomnia Symptom Questionnaire diagnostic threshold of insomnia. All had at worst mild symptoms of anxiety or depression, and most reported moderate consumption of alcohol and caffeine, moderate frequency of exercise, and only occasional naps. Six were current nicotine users, 14 reported using nicotine less than 100 times total, and 1 was a former nicotine user.

**Table 1.** Demographics of the sample. BMI: Body Mass Index; GAD : General Anxiety Disorder; ESS: Epworth Sleepiness Scale; PHQ-9 : Patient Health Questionnaire-9; ISI: Insomnia Sevrity Questionnaire

| | Mean ± SD | Min - Max |
| --- | --- | --- |
| female/male | 6/19 |  |
| Age | 35.32 ± 7.51 | 23 - 50 |
| BMI (kg.m-2) | 23.81 ± 3.43 | 17.44 - 31.6 |
| ISI | 5.00 ± 3.67 | 0 - 14 |
| ESS | 7.76 ± 3.77 | 1 - 19 |
| PHQ-9 | 1.84 ± 1.95 | 0 - 6 |
| GAD-7 | 2.00 ± 2.57 | 0 - 10 |
| N Naps/wk | 0.79 ± 1.13 | 0 - 4 |
| N Exercise/wk | 1.77 ± 1.72 | 0 - 6 |

### Protocol

Potential participants first completed a brief phone screen with study staff followed by an in-person interview at the French Armed Forces Biomedical Research Institute’s (IRBA) Fatigue and Vigilance Unit (Bretigny-Sur-Orge, France) during which they provided informed consent and subsequently completed a detailed demographic, medical, health, sleep, and lifestyle survey with a study physician to confirm eligibility. Once consented and eligibility was confirmed, participants were equipped by a sleep technologist to undergo an overnight sleep study at the center with simultaneous PSG and the DH recordings. The beginning and the end of the PSG and DH data collection periods were set based on participants’ self-selected lights-off and lights-on times. PSG and DH data recordings were synchronized a posteriori by resampling the DH data on the same timestamps as the PSG data so that records were perfectly aligned. Following the sleep study, technologists removed both devices, participants were debriefed and interviewed to identify any adverse events, and any technical problems were noted. All participants received financial compensation commensurate with the burden of study involvement. The study was approved by the Committees of Protection of Persons (CPP), declared to the French National Agency for Medicines and Health Products Safety, and carried out in compliance with the French Data Protection Act and International Conference on Harmonization (ICH) standards and the principles of the Declaration of Helsinki of 1964 as revised in 2013.

### Polysomnographic Assessment

The PSG assessment was performed using a Siesta 802 (Compumedics Limited, Victoria, Australia) with the following EEG derivations: F3/M2, F4/M1, C3/M2, C4/M1, O1/M2, O2/M1; 256 Hz sampling rate with a 0.03–35 Hz bandpass filter); bilateral electrooculographic (EOG), electrocardiographic (EKG), submental and bilateral leg electromyographic (EMG) recordings were also performed. Airflow, thoracic movements, snoring, and oxygen saturation were also monitored. EEG cup-electrodes of silver-silver chloride (Ag-AgCl) were attached to participants’ scalps with EC2 electrode cream (Grass Technologies, Astro-Med, Inc., West Warwick, RI, USA), according to the international 10-20 system for electrode placement. Auto-adhesive electrodes (Neuroline 720, Ambu A/S, Ballerup, Denmark) were used for EOG and EKG recordings.

### Study Device

The Dreem headband (DH) device is a wireless headband worn during sleep which records, stores, and automatically analyzes physiological data in real time without any connection (e.g., Bluetooth, Wi-Fi, etc.). Following the recording, the DH connects to a mobile device (e.g., smart phone, tablet) via Bluetooth to transfer aggregated metrics to a dedicated mobile application and via Wi-Fi to transfer raw data to the sponsor’s servers. Five types of physiological signals are recorded via 3 types of sensors embedded in the device: 1) brain cortical activity via 5 EEG dry electrodes yielding 7 derivations (FpZ-O1, FpZ-O2, FpZ-F7, F8-F7, F7-01, F8-O2, FpZ-F8; 250Hz with a 0.4-18 Hz bandpass filter); 2, 3, & 4) movements, position, and breathing frequency via a 3-D accelerometer located over the head; and 5) heart rate via a red-infrared pulse oximeter located in the frontal band (1). The EEG electrodes are made of high consistency silicone with soft, flexible protrusions on electrodes at the back of the head enabling them to acquire signal from the scalp through hair. An audio system delivering sounds via bone conduction transducers is integrated in the frontal band but was not assessed in this study. The DH is composed of foam and fabric with an elastic band behind the head making it adjustable such that it is tight enough to be secure, but loose enough to minimize discomfort. Additional details have been published previously in (19).

## Data Analysis

We divided data analysis into three parts: 1) EEG signal quality; 2) heart rate, breathing frequency, and RRV agreement; and 3) Automatic Sleep Stage classification of the DH compared to scorers’ consensus on the PSG.

### Assessing EEG Signal Quality

To assess the EEG signal quality of the DH, we computed the correlation of the relative spectral power in the *delta* (0.5 − 4*Hz*), *theta* (4 − 8*Hz*), *alpha* (8 − 14*Hz*), and beta (15 − 30*Hz*) bands during sleep throughout the night between the DH and the corresponding PSG records. Relative spectral power was computed on 90-second windows using a Fast Fourier Transform. Exponential smoothing with *alpha* = 0.7 was applied to the resulting signals to avoid abrupt transitions. The correlation was computed on the resulting smoothed signals. To maximize the time of the night with good signal quality on the DH, we developed a procedure to select a *virtual channel* which corresponds to the EEG frontal-occipital channel (FpZ-O1, FpZ-O2, F7-01, or F8-O2) with the best quality signal at any given epoch throughout the night; previously described in (19). We assessed spectral power on the virtual channel for the DH and compared it to a similar derivation on the PSG: F3-O1. As a baseline, we also compared the relative spectral power of the PSG F3-O1 PSG derivation to that of the F4-O1 PSG derivation for each frequency band using the same process. We excluded periods in which the *virtual channel* could not be computed on the DH signal because of bad signal quality on all channels (2.1% of the windows on average across all the recordings).

### Assessing Heart Rate, Breathing Frequency and Respiration Rate Variability agreement

The agreement of DH measurements of heart rate frequency (beats per minute) and breathing frequency (in cycle per minute) with PSG measurements of the same variables was also assessed. To do so, values were computed every 15 seconds (on 30-second sliding windows) on DH data and compared to the respective PSG values using an average of the Mean Absolute Error (MAE) computed for each record. An analogous method was employed to assess the capacity of the DH to retrieve respiration rate variability (RRV, in percentage), as described in (25).

#### Heart rate

Heart rate is recorded directly during a PSG. The DH, on the other hand, uses the pulse oximeter infrared signal to measure heart rate using a 3-step process:

1. Infrared signal is filtered between 0.4 and 2 Hz and zero crossing is applied to compute the mean heart rate frequency *fs*.
2. Infrared signal is filtered between *f s/*1.25 and *fs* ∗ 1.25 and zero crossing is applied to compute the minimum and maximum heart rate frequencies *fs*_*min*_ and *fs*_*max*_.
3. Infrared signal is filtered between *fs*_*min*_ and *fs*_*max*_, zero crossing is applied and heart rate is computed on 15-second windows. The heart rate measured by the DH was compared to each respective 15-second window obtained from the PSG. Exponential smoothing with *alpha* = 0.3 was applied on both the DH and PSG heart rates to avoid brutal transitions.

This standard method provides a robust measure of heart rate frequency during sleep. However, it would probably be ill-suited for waking measurement where artefacts and noise are more likely to occur. One record was excluded from heart rate analyses because the PSG heart rate measurement remained at the same value for the entire duration of the record, and was therefore assumed to be inaccurate.

#### Breathing Frequency

Breathing frequency was measured by the z-axis of the accelerometer on the DH and by the external pressure signal of the PSG. To compute breathing frequency from the DH signals when the participant was asleep, an analogous 3-step process to that for the heart rate computation was followed, using a filter between 0.16 and 0.3 Hz in the first step.

#### Respiration Rate Variability

The RRV is computed with the exact same methodology than the one described in (25), except that the method employed here computed RRV throughout the entire night instead of on steady sleep windows. For the PSG, the method was applied to the external pressure channel. For the DH, the RRV was computed on the 3 axes of the accelerometer and the minimum value between the three was kept for each computed value to reduce noise.

### Assessing Sleep Stages Classification Performance

Due to known inter-rater variance among even expert sleep scorers (26), using a single rater as the reference point renders comparison vulnerable to unintended bias. Thus, each PSG records was independently scored by 5 trained and experienced registered sleep technologists from 3 different sleep centers following the guidelines of the AASM (4). The DH data was scored by the embedded automatic algorithm of the DH.

#### Scoring performance metrics

To measure agreement between two hypnograms on a record, *Accuracy* (ratio of correct answers) and *Cohen’s Kappa*, 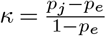 where *p*_*j*_ is the scorer accuracy and *p*_*e*_ is the baseline accuracy, are provided. We also computed the *F1-score* because it takes into account both *Precision* and *Recall*, as well as class imbalance, making it a rigorous metric for evaluating performance (27).

It is computed as: 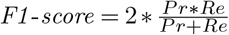 with 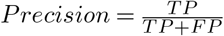 and 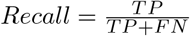, where TP, FP, and FN are the number of true positives, false positives, and false negatives, respectively. This score is computed per-class and averaged taking the weight of each class into account.

For ‘overall’ analyses, the average of the respective values from each individual record is calculated.

#### Scoring performance metrics evaluation

To evaluate scoring performance metrics and benefit from the multiple sleep experts scorings, a similar methodology to (12) was used. Indeed, to evaluate the performance metrics for each scorer, the scoring from each individual scorer was compared to the consensus scoring of the four other scorers. To evaluate the performance metrics of the DH automatic approach, the automatic scoring from the DH was compared to the consensus scoring of the four top-ranked scorers. This method ensures that both the individual scorers and the automatic algorithm running on the DH data were evaluated against a consensus of exactly four scorers.

#### Building a consensus scoring from multiple scorings

Thus, we developed a way to build a unique consensus scoring from multiple scorings on a record. For each epoch, the majority opinion across scorers is chosen. In case of a tie, the sleep stage scored by the top ranked scorer is used (scorer ranking procedure described below, as Soft-Agreement); ties occurred on 7.3 ± 2.4% of the epochs on average across all the records.

#### Scorer ranking

The previous section highlights the need to rank scorers in order to build a valid consensus scoring. The ranking of a scorer is based on his level of agreement with all the other scorers. To measure this, we introduce below an agreement metric between one scoring against multiple other scorings. We call this metric ‘Soft-Agreement’ as it takes all the scorings into account and does not require any thresholding.

##### Notations

Let *y*_*j*_ ∈ ⟦4⟧^*T*^ be the sleep staging associated to scorer *j* taking values in {0, 1, 2, 3, 4} standing respectively for Wake, N1, N2, N3 and REM with size *T* epochs. Let *N* be the number of scorers. Let 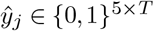 be the one hot encoding of *y*_*j*_. For each epoch *t* ∈ ⟦*T*⟧ its value is 1 for the scored stage and 0 for the other stages.

First, we define a probabilistic consensus 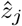 as:

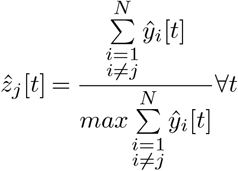

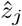 takes values in [0, 1]^5×*T*^. For each epoch *t* ∈ ⟦*T*⟧, the value for each sleep stage is proportional to the number of scorers who scored that sleep stage. A value of ‘1’ is assigned if the chosen stage matches the majority sleep stage or any of the sleep stages involved in a majority tie. We then define the Soft-Agreement for scorer *j* as:

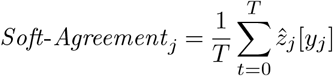

A Soft-Agreement of 1 means that for all epochs, scorer *j* scored the same sleep stage as the majority and, in case of tie, he scored one of the sleep stage involved in the tie. A Soft-Agreement of 0 would happen if scorer *j* systematically scores a different stage than *all* of the other scorers.

To rank the five scorers in this study, the Soft-Agreement was computed for each scorer against the four others on each record and then averaged across all the records. Based on these values, we are able to build unique consensus scorings for comparison with each scorer. To build the consensus scorings for comparison with the DH automatic algorithm, scorings from the top-four scorers were used.

#### Performance assessment of sleep variables

The following standard sleep variables were calculated: time in bed (TIB), as the number of minutes from lights-out to lights-on; total sleep time (TST, min); sleep efficiency (SE, %), as TST / TIB * 100; sleep onset latency (SOL), as the number of minutes from lights-out to the first three consecutive epochs of any sleep stage; wake after sleep onset (WASO), as the number of minutes awake following the first three consecutive epochs of any sleep stage; and the time (min) and percentage of TST spent in each sleep stage (N1, N2, N3, and REM).

#### Dreem Headband Algorithm

The DH embedded automatic algorithm works in 2 stages: 1) feature extraction and 2) classification. It is able to provide real-time sleep staging predictions. 1) Feature extraction is performed for each new epoch of 30 seconds. Features extracted from the various sensors are concatenated to go through the classification layer. EEG features include power frequency in the delta, alpha, theta, and beta bands and ratio of relative powers as described in (23). Sleep patterns (e.g. slow oscillations, alpha rhythm, spindles, K-complexes) are detected using an expert approach. The accelerometer provides breathing, movement, and position features. The pulse oximeter provides cardiac features. A total of 79 features are extracted from each raw DH record. 2) The classification module is built from two layers of Long-Short Term Memory (28) and a Softmax function outputting the final probability prediction that the epoch belongs to each sleep stage. It relies on the features extracted from the last 30 epochs to predict the current one. Hence, it takes into account the past temporal context to make a prediction, as a sleep expert would do. This classification module is trained using backpropagation. The training has been done on a dataset composed of previously recorded internal Dreem records. A total of 423 records were used for training and presented several times to the network. 213 validation records from other subjects were used to stop the training when the performance metrics computed on this validation set were not improving anymore. None of the records of the current study were used to train or validate the network. We used the framework provided by Pytorch (29) and trained on a single Nvidia Titan X GPU (~1 hour of training, ~1 seconds for inference).

## Results

The data collection was well tolerated with no adverse effects reported for the DH. The set up time was ~5 min for the DH and ~45 min for the PSG.

### EEG Signal Quality

The quality of the EEG signal assessed through the correlation of the relative spectral power between DH and PSG for alpha, beta, delta, and theta frequencies is presented in the Table 2. Results indicate a substantial agreement between the DH and PSG derivations for all of the frequency bands (> 0.6, p-values were < 1e-10). As expected, the correlations between the DH and the PSG are lower than the baseline correlations between the 2 derivations from the same PSG record (F3-O1 and F4-O2). The Figure 2 shows a sample of raw signals recorded by the DH and a PSG on the same record during each sleep stage (N1, N2, N3, REM, Wake). Figure 3 shows the relative spectral power between DH and PSG for each EEG frequency examined throughout a representative record.

**Table 2.** Pearson correlation for alpha, beta, delta and theta EEG frequency for the Dreem headband (DH) vs PSG1 (F3-O1) and PSG1 (F3-O1) vs PSG2 (F4-O2).

|  | DH / PSG F3-O1 | PSG F3-O1 / PSG F4-O2 |
| --- | --- | --- |
| alpha | $0.71 \pm 0.13$ | $0.85 \pm 0.12$ |
| beta | $0.71 \pm 0.18$ | $0.91 \pm 0.08$ |
| delta | $0.76 \pm 0.14$ | $0.90 \pm 0.06$ |
| theta | $0.61 \pm 0.12$ | $0.82 \pm 0.07$ |

**Fig. 1.**
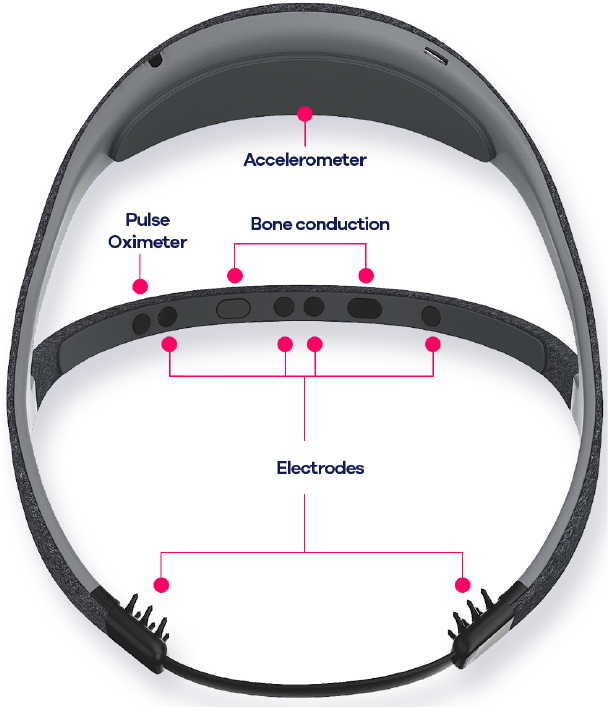
Illustration of the Dreem headband (DH) presenting the location of the various sensors. The top arch contains the accelerometer, the battery, and the electronics. The front band contains all of the other sensors and the audio bone conduction except for the two electrodes positioned at the back of the head.

**Fig. 2.**
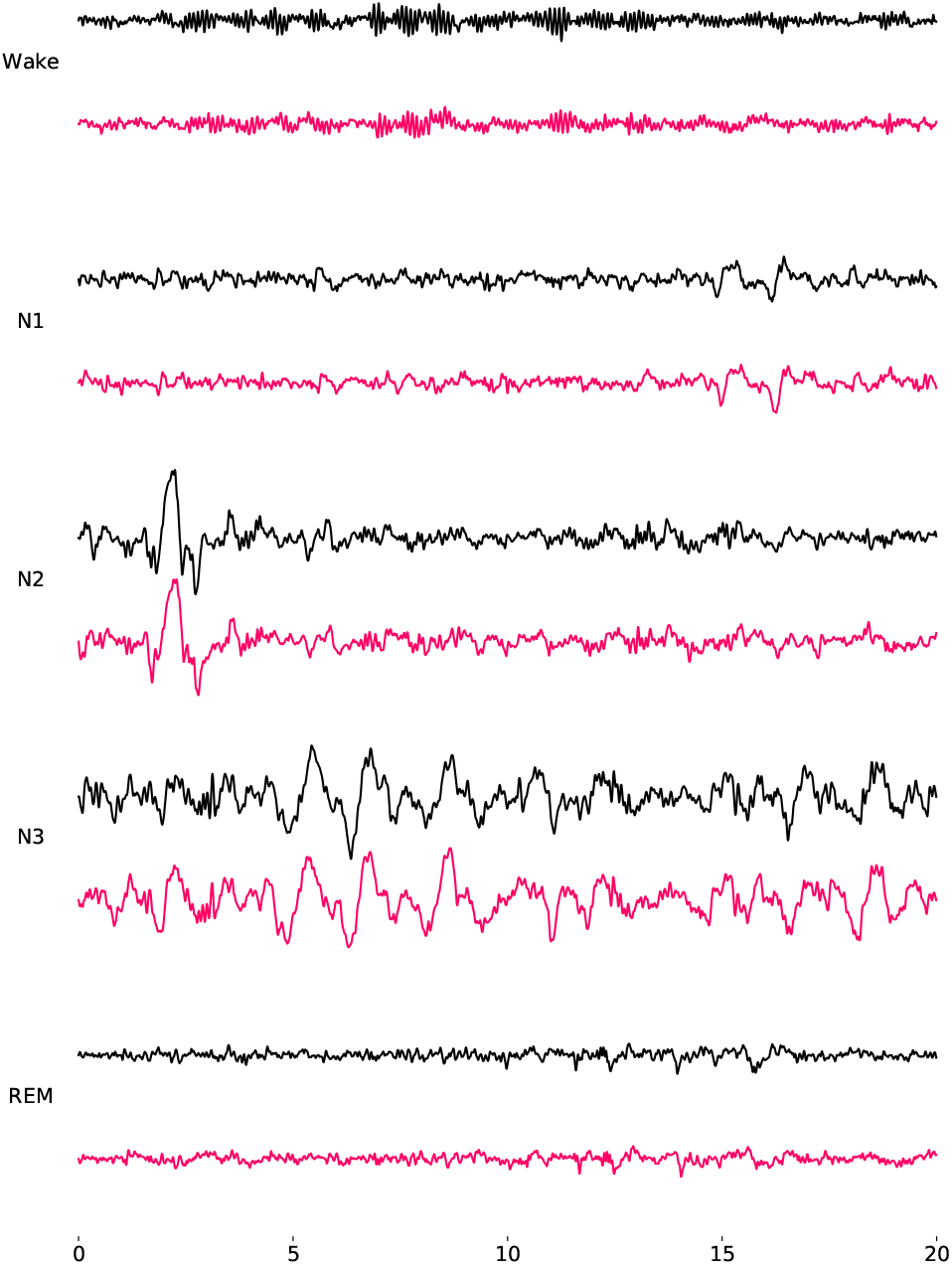
20-second samples of raw signals recorded by Dreem headband (DH, pink) and polysomnography (PSG, black) on the same record during each sleep stages (N1, N2, N3, REM, Wake). The derivations are F7-O1 for the DH and F3-O1 for the PSG. The signals are presented between −150 and 150 uV.

**Fig. 3.**
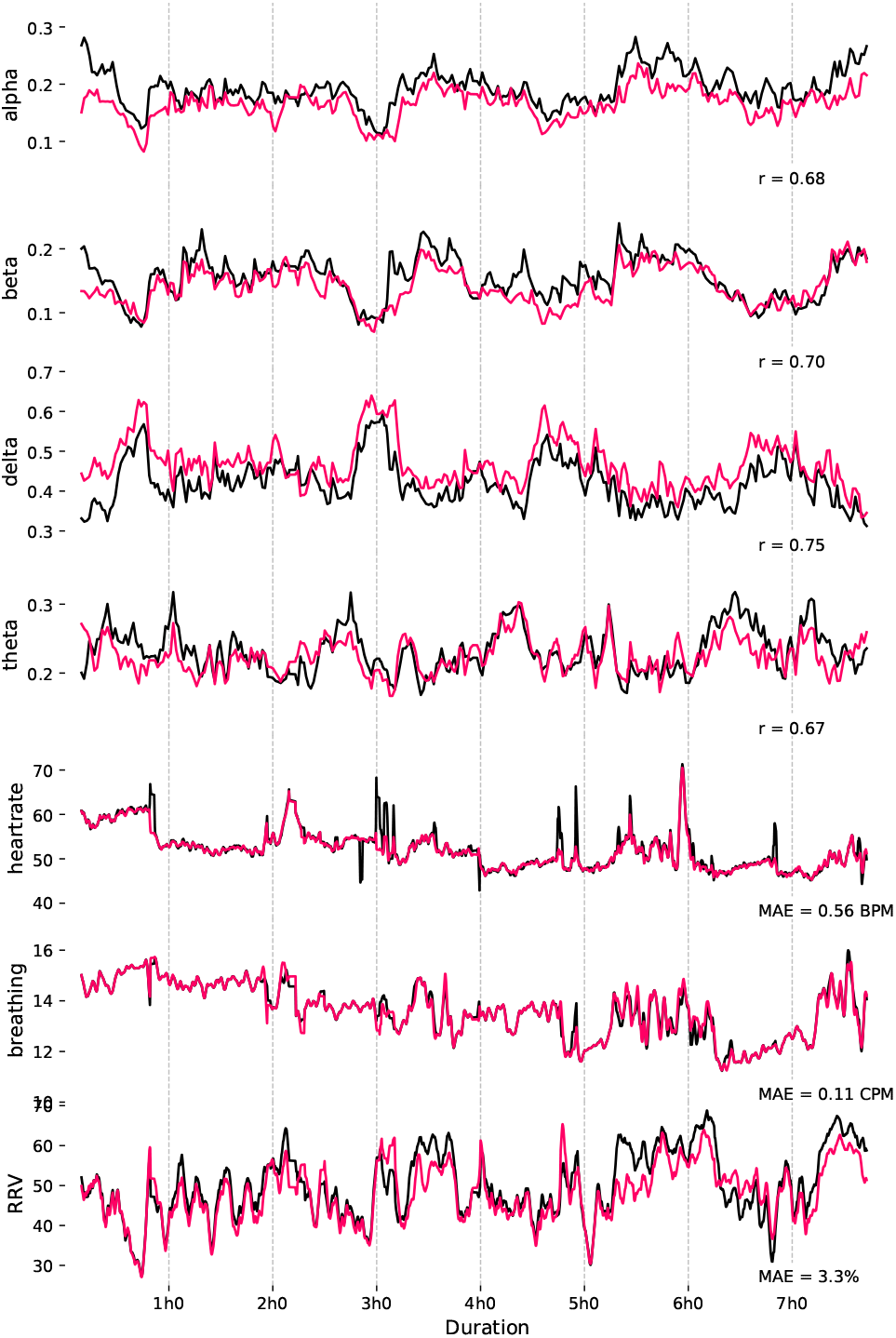
Relative spectral power (alpha, beta, delta, and theta frequencies, A.U.), heart rate (beats per minute, BPM), breathing frequency (cycles per minute, CPM), and respiratory rate variability (RRV, %) for a representative record (i.e., with a correlation value similar to the mean of the group). These signals are presented for a whole record for both the Dreem headband (DH, pink) and polysomnography (PSG, black).

### Heart Rate and Breathing Agreement

The agreements of heart rate, breathing frequency, and respiratory rate variability measured by the DH and those measured by the PSG are presented in Table 3. The results show an excellent agreement of the DH compared to the PSG with minimum absolute errors of 1.2 ± 0.5 bpm, 0.3 ± 0.2 cpm, and 3.2 ± 0.6% for the heart rate, breathing frequency, and RRV during sleep, respectively. Figure 3 shows an example of heart rate, breathing frequency, and RRV measured on the PSG throughout the night.

**Table 3.**
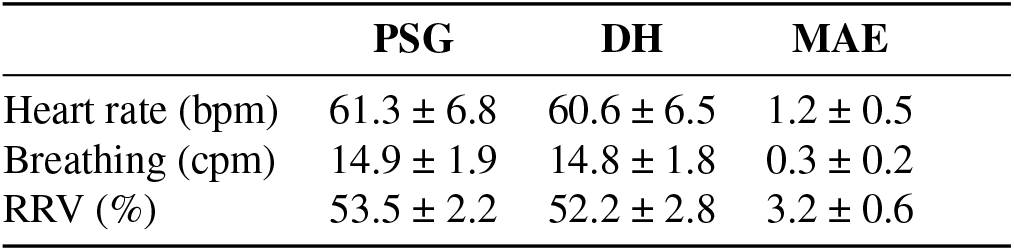
The Dreem headband (DH) and Polysomnography (PSG) columns presents mean values ± SD computed across all records for heart rate, breathing frequency and respiratory rate variability (RRV) for both devices. Average mean absolute error (MAE) by record is given in the last column.

### Sleep Stage Classification

The five scorers had Soft-Agreement scores of 88.6%, 90.7%, 91.7%, 84.2% and % for scorers 1, 2, 3, 4 and 5, respectively, resulting in an overall Soft-Agreement score of 89.4 ± 2.79%.

With these Soft-Agreement values, we were able to develop consensuses with which to compare each scorer and the predictions of the DH automatic algorithm for the purpose of evaluating the metrics presented in Table 4. The overall accuracy of the five scorers was 86.4 ± 7.4%, with scorer 1 = 86.3 ± 10.5%, scorer 2 = 88.2 ± 4.2%, scorer 3 = 88.9 ± 5.1%, scorer 4 = 82.0 ± 8.1%, scorer 5 = 88.9 ± 4.6%. Notably, these accuracies are above the average performance of other certified scorers reported in the literature (8), indicating the scorers in this study were well-trained. Across the manual scorers, accuracy was highest for REM sleep (87.8 ± 13.6%) and followed closely by N2 (85.9 ± 10.7%) and N3 sleep stages (84.2 ± 20.6%). The accuracy for wake was slightly lower (82.5 ± 17.5%) and lowest for N1 (54.2 ± 16.8%). Variability among the scorers was largest for the N3 sleep stage with a standard deviation of 20.6%.

**Table 4.** Performance metrics for each scorer and the automatic approach of the Dreem headband (DH) computed by comparison against their consensus. Overall column presents Mean ± SD observed for the five scorers. Results are given for each sleep stages.

|  |  | DH | Overall Scorers | Scorer 1 | Scorer 2 | Scorer 3 | Scorer 4 | Scorer 5 |
| --- | --- | --- | --- | --- | --- | --- | --- | --- |
| **All** | F1 (%) | 83.8 ± 6.3 | 86.8 ± 7.4 | 86.3 ± 10.5 | 88.2 ± 4.2 | 88.9 ± 5.1 | 82.0 ± 8.1 | 88.9 ± 4.6 |
|  | Accuracy (%) | 83.5 ± 6.4 | 86.4 ± 8.0 | 85.7 ± 12.1 | 87.5 ± 4.5 | 88.9 ± 4.6 | 81.2 ± 8.8 | 88.9 ± 4.2 |
|  | Cohen Kappa (%) | 74.8 ± 10.4 | 79.8 ± 11.4 | 78.9 ± 15.7 | 81.2 ± 7.0 | 83.2 ± 7.2 | 72.5 ± 13.2 | 83.0 ± 7.2 |
| **wake** | F1 (%) | 76.7 ± 14.3 | 84.1 ± 13.6 | 85.9 ± 10.1 | 86.5 ± 12.3 | 87.6 ± 9.9 | 74.9 ± 18.0 | 85.6 ± 11.9 |
|  | Accuracy (%) | 74.0 ± 18.1 | 82.5 ± 17.5 | 80.2 ± 14.0 | 78.1 ± 19.3 | 90.2 ± 12.9 | 82.4 ± 21.0 | 81.6 ± 16.5 |
|  | Cohen Kappa (%) | 74.1 ± 15.2 | 82.2 ± 15.0 | 84.4 ± 10.7 | 85.2 ± 12.6 | 86.3 ± 10.4 | 71.1 ± 20.5 | 84.1 ± 12.8 |
| **N1** | F1 (%) | 46.5 ± 12.4 | 49.7 ± 14.5 | 49.3 ± 13.8 | 51.2 ± 11.4 | 53.7 ± 13.7 | 39.7 ± 16.3 | 54.5 ± 11.6 |
|  | Accuracy (%) | 47.7 ± 15.6 | 54.2 ± 16.8 | 58.2 ± 14.7 | 60.3 ± 12.6 | 59.1 ± 14.4 | 38.3 ± 15.5 | 54.9 ± 16.3 |
|  | Cohen Kappa (%) | 43.5 ± 12.6 | 47.0 ± 15.1 | 46.6 ± 14.0 | 48.5 ± 11.8 | 51.4 ± 13.9 | 36.4 ± 17.3 | 52.3 ± 12.1 |
| **N2** | F1 (%) | 87.5 ± 5.5 | 89.0 ± 7.3 | 88.2 ± 11.8 | 90.3 ± 4.6 | 90.7 ± 4.1 | 84.3 ± 6.9 | 91.3 ± 3.5 |
|  | Accuracy (%) | 82.9 ± 8.1 | 85.9 ± 10.7 | 87.8 ± 13.6 | 87.6 ± 8.7 | 89.3 ± 6.0 | 75.8 ± 10.9 | 89.0 ± 5.7 |
|  | Cohen Kappa (%) | 75.4 ± 10.8 | 78.8 ± 13.2 | 77.5 ± 20.5 | 81.1 ± 8.6 | 81.5 ± 8.5 | 71.1 ± 12.9 | 82.9 ± 7.1 |
| **N3** | F1 (%) | 76.4 ± 22.9 | 78.3 ± 23.8 | 81.0 ± 24.1 | 79.6 ± 22.8 | 76.8 ± 25.0 | 75.2 ± 22.1 | 78.9 ± 24.7 |
|  | Accuracy (%) | 82.6 ± 20.6 | 84.2 ± 20.6 | 89.1 ± 14.0 | 89.1 ± 11.1 | 66.9 ± 27.6 | 92.7 ± 12.0 | 84.1 ± 21.3 |
|  | Cohen Kappa (%) | 74.0 ± 22.6 | 76.6 ± 23.1 | 79.2 ± 24.1 | 77.2 ± 22.8 | 74.8 ± 25.0 | 74.9 ± 17.9 | 76.8 ± 24.3 |
| **REM** | F1 (%) | 82.9 ± 12.3 | 90.9 ± 10.2 | 85.0 ± 17.0 | 91.4 ± 4.2 | 94.4 ± 4.6 | 91.5 ± 8.8 | 92.1 ± 8.1 |
|  | Accuracy (%) | 84.5 ± 13.3 | 87.8 ± 13.6 | 76.3 ± 19.4 | 85.8 ± 6.5 | 92.1 ± 9.5 | 89.5 ± 13.6 | 95.3 ± 3.1 |
|  | Cohen Kappa (%) | 79.0 ± 13.7 | 89.0 ± 11.1 | 82.5 ± 17.6 | 89.4 ± 5.3 | 93.1 ± 5.1 | 89.7 ± 10.5 | 90.5 ± 8.9 |

The overall accuracies of the DH automated algorithm using the DH data for sleep staging compared to the scorer consensuses are presented in Table 4 (overall accuracy = 83.8 ± 6.8%). The classification accuracy per stage of the DH parallels the order of the manual scorers using PSG data: highest accuracy for REM sleep (84.5 ± 13.3%) followed by N2 (82.9 ± 8.1%) and N3 sleep stages (82.6 ± 20.6%). The accuracy for wake was lower (74.0 ± 18.1%) with the lowest accuracy similarly obtained for the N1 sleep stage (47.7 ± 15.6%). The confusion matrices (Figure 4) shows the classifications per stage of both the DH and scorer averages versus the respective consensuses. According to the matrices, both in the case of the DH and the PSG scoring, Wake was most often misclassified as N1 (12.4% a nd 1 0.5% o f e pochs, respectively), and N3 was most often misclassified as N 2 (16.4% and 20.5% of epochs for DH and PSG, respectively). Figure 6 shows five representative hypnograms computed from the DH classifications a nd t he c orresponding s corer consensus hypnograms.

**Fig. 4.**
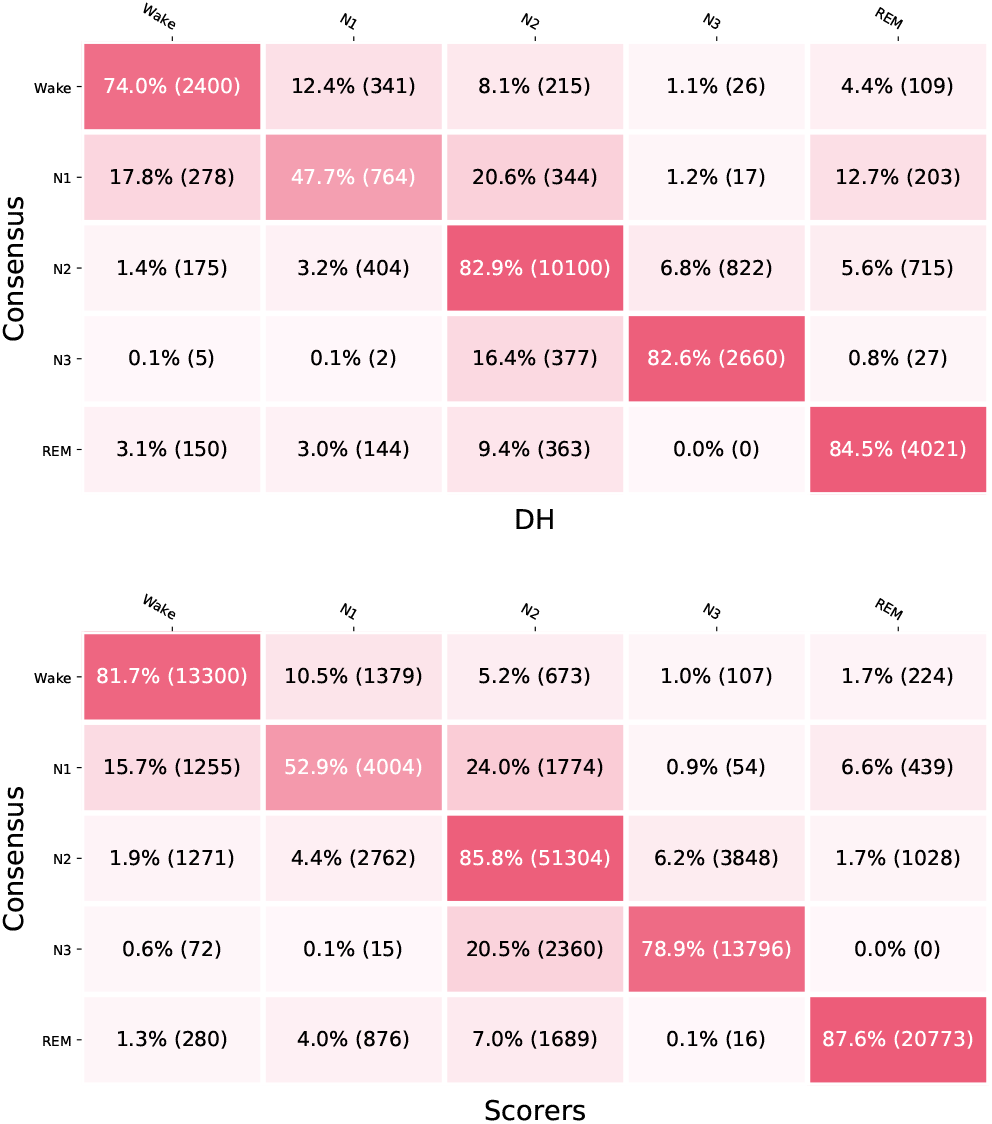
Confusion matrix for the Dreem headband (DH) versus PSG scoring consensuses (top) and the overall confusion matrix for scorers versus the other scorers’ consensuses (bottom). Values are normalized by row with the number of epochs in parentheses.

Table 5 reports analyses of sleep variables traditionally used in sleep medicine computed for the DH and scorers’ averages versus scorers’ consensuses, as well as differential results for the DH and overall differentials for the scorers. Results indicate high similarity between the DH measurements and the consensuses and a similar level of variability for the DH and the average scorers’ values. Table 6 shows a high level of correlation of each sleep variable with the scorers’ consensus values. Bland Altman plots are presented in Figure 5 and show high agreement between the DH and the consensus scores, low bias, low fluctuation around the mean, and no clear outliers, for all of the sleep variables analyzed.

**Table 5.** The consensus column presents sleep variables computed with the scorers consensus of the top-four ranked scorers. The Dreem headband (DH) column present the sleep variables computed on the DH. Differential DH column presents the average per-record difference observed between the DH and the scorer consensus. Overall Differential Scorers presents the average per-record difference observed between each scorer and the scorer consensus formed by the four other scorers. Results are presented as Mean ± SD [0.95CI].

|  | Consensus | DH | Differential DH | Overall Differential Scorers |
| --- | --- | --- | --- | --- |
| TST (min) | 430.4 ± 66.8 [402.2, 458.5] | 431.8 ± 68.4 [402.9, 460.6] | 1.4 ± 19.0 [-6.6, 9.4] | -2.0 ± 28.5 [-7.1, 3.0] |
| Sleep efficiency (%) | 87.0 ± 8.1 [83.5, 90.4] | 87.3 ± 8.4 [83.8, 90.8] | 0.3 ± 4.0 [-1.4, 2.0] | -0.4 ± 5.6 [-1.4, 0.6] |
| SOL (min) | 18.9 ± 18.7 [11.0, 26.8] | 20.0 ± 18.2 [12.4, 27.7] | 1.2 ± 3.6 [-0.4, 2.7] | -1.7 ± 15.6 [-4.5, 1.1] |
| WASO (min) | 44.1 ± 27.8 [32.4, 55.8] | 41.5 ± 31.7 [28.1, 54.9] | -2.6 ± 19.5 [-10.8, 5.7] | 3.7 ± 28.4 [-1.3, 8.8] |
| Stage Wake duration (min) | 61.8 ± 38.8 [45.5, 78.1] | 60.2 ± 40.7 [43.0, 77.3] | -1.7 ± 19.1 [-9.7, 6.4] | 2.0 ± 28.5 [-3.1, 7.0] |
| Stage Wake (%) | 12.8 ± 8.1 [9.4, 16.2] | 12.4 ± 8.3 [8.9, 15.9] | -0.4 ± 4.0 [-2.1, 1.3] | 0.4 ± 5.6 [-0.6, 1.4] |
| Stage N1 duration (min) | 31.0 ± 14.4 [25.0, 37.1] | 31.9 ± 17.9 [24.4, 39.4] | 0.8 ± 12.9 [-4.6, 6.3] | 6.0 ± 25.2 [1.5, 10.5] |
| Stage N1 (%) | 6.3 ± 2.9 [5.1, 7.5] | 6.5 ± 3.7 [5.0, 8.1] | 0.2 ± 2.6 [-0.9, 1.3] | 1.2 ± 4.4 [0.4, 1.9] |
| Stage N2 duration (min) | 244.3 ± 59.2 [219.4, 269.2] | 227.9 ± 57.0 [203.9, 251.9] | -16.4 ± 21.8 [-25.5, -7.2] | -9.7 ± 38.7 [-16.6, -2.8] |
| Stage N2 (%) | 49.2 ± 9.3 [45.3, 53.1] | 46.1 ± 9.9 [41.9, 50.2] | -3.1 ± 4.3 [-5.0, -1.3] | -1.8 ± 7.5 [-3.2, -0.5] |
| Stage N3 duration (min) | 61.4 ± 30.4 [48.6, 74.2] | 70.5 ± 37.1 [54.8, 86.1] | 9.1 ± 19.5 [0.8, 17.3] | 6.3 ± 32.8 [0.5, 12.1] |
| Stage N3 (%) | 12.5 ± 6.1 [9.9, 15.0] | 14.2 ± 6.9 [11.3, 17.1] | 1.7 ± 3.8 [0.1, 3.3] | 1.3 ± 6.1 [0.2, 2.3] |
| REM sleep duration (min) | 93.6 ± 32.3 [80.0, 107.2] | 101.5 ± 41.7 [83.9, 119.1] | 7.9 ± 24.9 [-2.5, 18.4] | -4.7 ± 16.7 [-7.7, -1.7] |
| REM sleep (%) | 19.0 ± 6.2 [16.4, 21.6] | 20.5 ± 7.9 [17.1, 23.8] | 1.5 ± 5.0 [-0.6, 3.6] | -1.0 ± 3.3 [-1.5, -0.4] |

**Table 6.** Pearson correlation of the sleep parameters computed on every recordings with the corresponding p-value.

|  | Correlation Coef. | p-Value |
| --- | --- | --- |
| TST (min) | 0.96 | 2.8e-14 |
| Sleep efficiency (%) | 0.89 | 4.2e-09 |
| SOL (min) | 0.98 | 7.3e-18 |
| WASO (min) | 0.79 | 2.2e-06 |
| Stage Wake duration (min) | 0.89 | 4.0e-09 |
| Stage Wake (%) | 0.88 | 5.7e-09 |
| Stage N1 duration (min) | 0.7 | 9.7e-05 |
| Stage N1 (%) | 0.71 | 8.1e-05 |
| Stage N2 duration (min) | 0.93 | 1.6e-11 |
| Stage N2 (%) | 0.9 | 9.6e-10 |
| Stage N3 duration (min) | 0.85 | 6.8e-08 |
| Stage N3 (%) | 0.84 | 1.9e-07 |
| REM sleep duration (min) | 0.8 | 1.4e-06 |
| REM sleep (%) | 0.78 | 4.2e-06 |

**Fig. 5.**
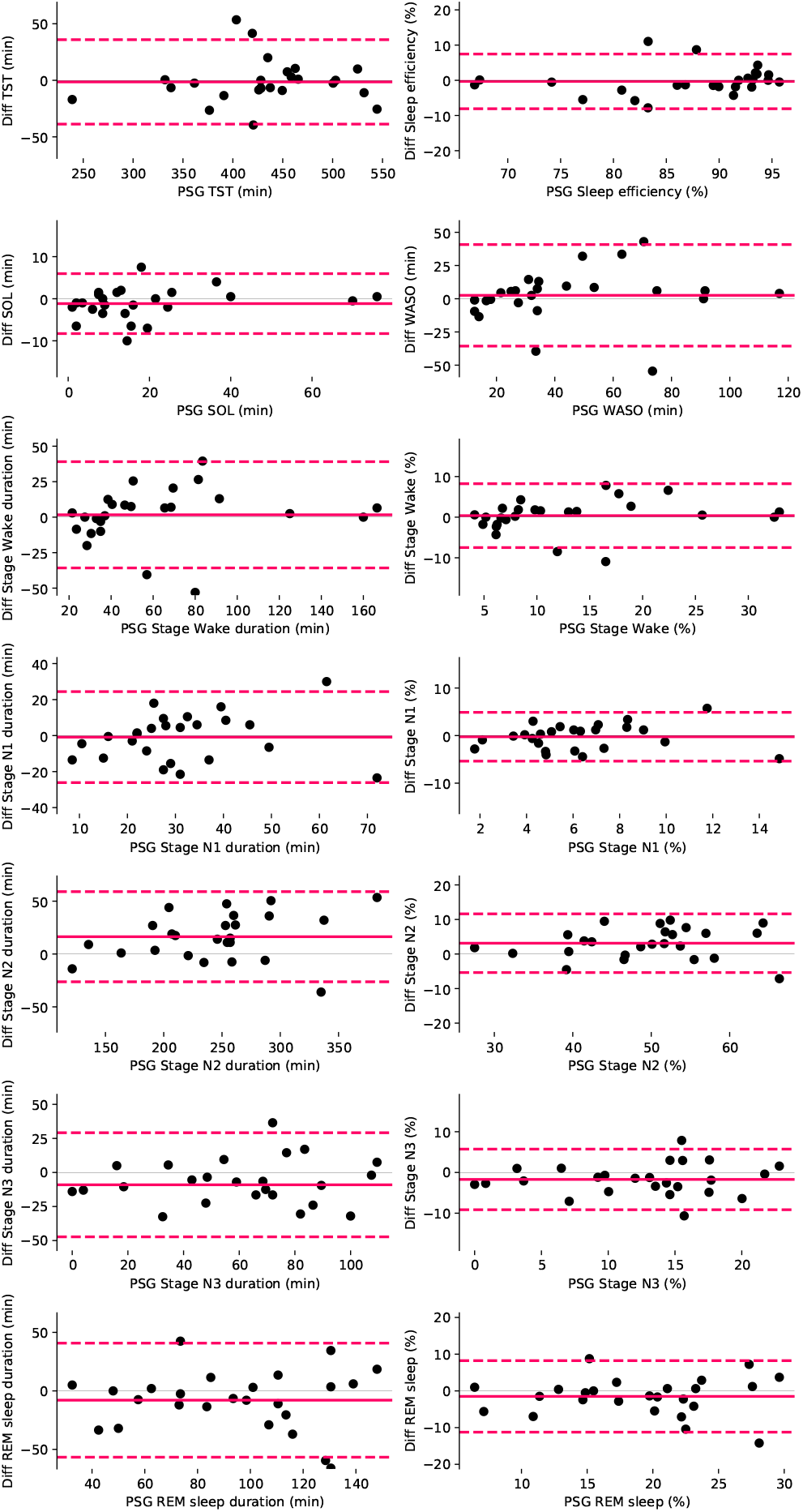
Bland Altman plots for each sleep variable measures by the Dreem head-band (DH) versus the consensus sleep metrics computed for each record.

**Fig. 6.**
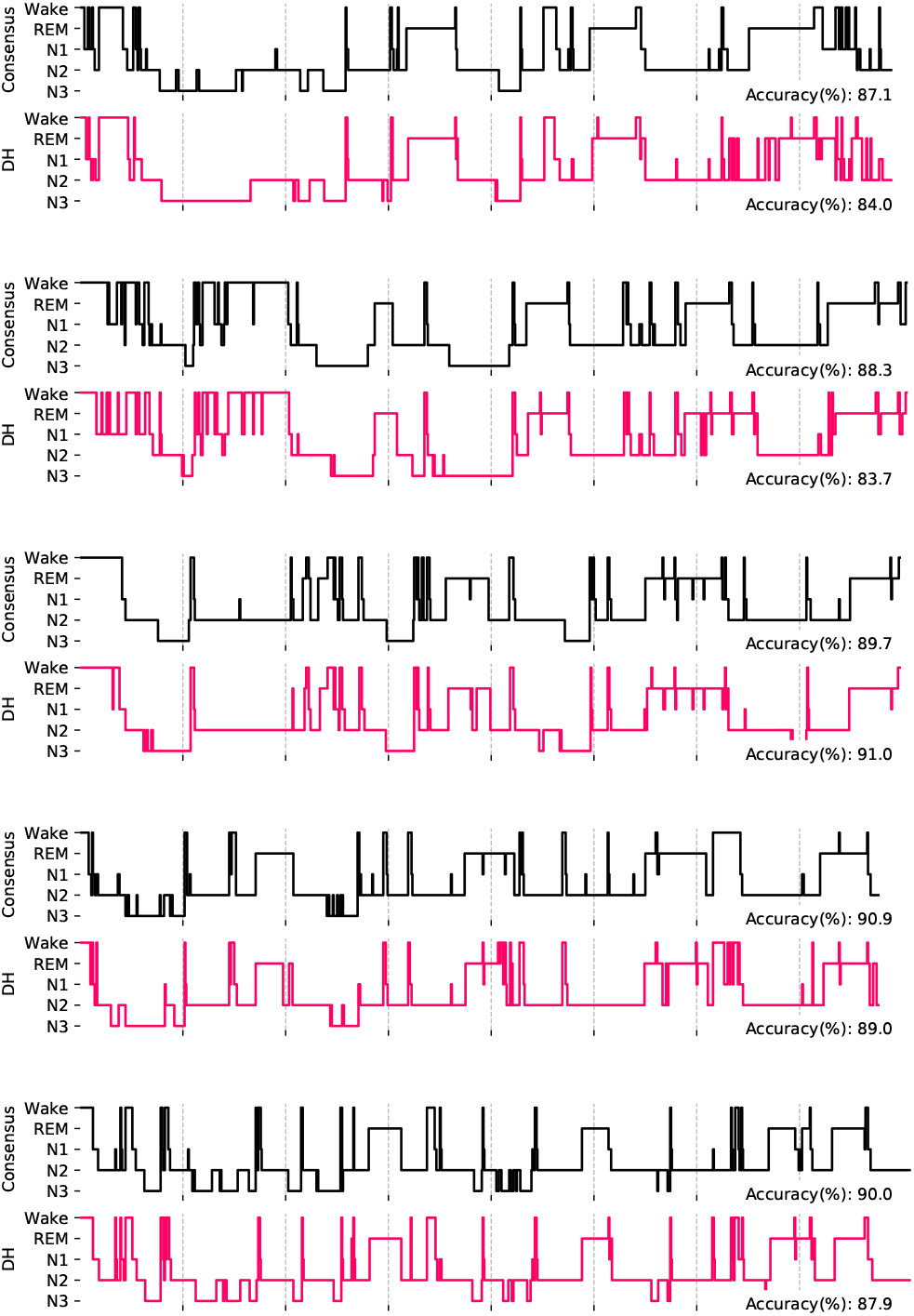
Hypnograms for the five first participants showing both the consensuses of the four top-ranked scorers (gray) and the DH automated sleep stage classifications. Accuracies are presented as average obtained by the five scorers on the consensus hypnogram, and scores obtained for the DH versus the consensuses.

## Discussion

This study compared the EEG signal quality; heart rate, breathing frequency, and respiratory rate variability (RRV) measurements; and automatic sleep staging of the Dreem headband (DH) to manually generated consensus scores of simultaneous in-lab PSG recordings. The data demonstrate that the DH: 1) acquires EEG during sleep with signal quality sufficient to enable reliable EEG-based sleep studies; 2) reliably measures breathing frequency and heart rate continuously during sleep; and 3) can perform automatic sleep staging classification according to AASM c riteria with performance similar to that of a consensus of 5 scorers using medical-grade PSG data.

First, we showed that EEG frequencies traditionally used in sleep medicine can be measured using dry electrodes with a substantial agreement with a PSG (30) despite the fact that the signal compared was not on the exact same derivation. The similar results in other domains comparing wet and dry electrodes have shown that these technologies are comparable for monitoring EEG signals paving the way to meaningful physiological monitoring at home under various conditions (31, 32). The levels of correlation between the DH and the PSG scores allows a trained human to identify typical sleep EEG patterns such as alpha rhythm, spindles, delta waves, sawtooth waves, or k-complexes. Recording raw physiological signals is critical for sleep research because sleep stages provide only limited insight into sleep quality. Furthermore, these specific patterns can now be automatically detected using deep neural networks (15, 33); though, analyses of the latter patterns are not reported here.

Second, the data show that our method for detecting breathing frequency and RRV using an accelerometer has excellent agreement with the gold standard. The position of the 3-D accelerometer, located over the head, appears to be a sensitive location for detecting small movements. These breathing measures are of interest during sleep because they indicate sympathovagal tone and potential sleep apnea syndrome. The excellent agreement for heart rate is similar to other studies showing that an infrared pulse oximeter positioned against the forehead can be used to reliably monitor heart rate. However, we were unable to provide a heart rate variability on most of the records due to insufficient resolution, similar to other studies (34). Further investigation of the DH in clinical populations with a suspicion of sleep apnea should be conducted.

Third, we showed that the DH is able to perform real-time sleep staging using data collected by the DH with an accuracy in the range of individual scorers using PSG data and comparable to the accuracy between PSG scorers in other studies (8, 26). To our knowledge, this performance on a dry EEG wearable has never been achieved with another device. All of the sleep variables computed by the DH are highly correlated with the expert consensuses with the highest correlation coefficient for TST (r = 0.96) and the lowest for the N1 sleep stage (r = 0.7). Sleep variables are macro-metrics computed on the hypnogram and are less impacted than sleep staging metrics by local differences. For instance, wake is slightly underestimated but that does not significantly impact sleep variables related to wake (WASO, sleep latency, sleep efficiency). Even though the inter-scorer reliability achieved with PSG by our 5 scorers was high, it highlights the need for such validation studies to rely on a consensus of multiple sleep experts when analyzing sleep staging performance (26). Mixing sleep experts from different sleep centers provides a more realistic analysis than is typically obtained in a standard clinical sleep study where records are scored by only a single individual, which strengthens our results. To evaluate these individual scorers, we introduced an objective methodology to build a consensus from the other scorers. This enables a fair evaluation of both individual scorers and the automated algorithmic approach of the DH. Interestingly, lower inter-scorer performance correlates with lower performance on the DH (data not shown), further reinforcing the similarities between the DH and manually scored PSG.

The main limitation of this study is that the sample was somewhat small and homogeneous in age and sleeper profile; even though this is consistent with the majority of similar validation studies (20, 22, 23). A larger sample of more diverse sleepers would have provided more reliability and generalizability to the general population. Notably, we excluded 2.1% of the windows on average across all the recordings in which the *virtual channel* could not be computed on the DH signal because of bad signal quality on every channel. Importantly too, the study includes only one night of data per subject. This remains a potential limitation due to lack of habituation to sleeping with a full PSG in a clinical sleep lab, which often leads to sleep being shorter and more fragmented; although our sample did achieve 87% sleep efficiency on average, suggesting that sleep was not substantially disrupted on a wide scale in this study. Thus, a single night of PSG data from a sleep lab may not be a reliable representation of typical sleep in the natural home environment.

## Conclusion

Taken together, the results of our analyses strongly support the utility of the Dreem headband for reliably recording high-quality sleep EEG, heart rate, and breathing with clinical-level performance of real-time automatic sleep analysis. While considerable value could be derived from longitudinal sleep EEG monitoring, until recently, (wet) EEG electrodes were too impractical to be easily used on a regular basis and without assistance. Our dry electrodes are considerably more comfortable and easier to apply without substantially sacrificing signal acquisition quality. Indeed, EEG pattern detection (alpha, spindles, K-complexes) could be performed by an automatic approach running on dry electrodes such as an unlock large scale analyses and studies of those pattern characteristics (15). Moreover, the Dreem head-band is able to measure breathing, and such pattern detection methods could be adapted to detect breathing events linked to apnea. The analysis described in this study has been developed to be used in a real-time setup to allow the Dreem head-band to support biofeedback and neurofeedback-based applications (19). These results, together with the price, ease of use, precision and reliability, and the collection of raw EEG and other relevant physiological data, make the Dreem head-band an ideal candidate for high-quality large-scale longitudinal sleep studies in the home or laboratory environment. As such, this technology can enable groundbreaking advancements in sleep research and medicine. For instance, the resulting database can ultimately be integrated with other types of data collection devices and used to identify unknown patient subgroups, detect early disease biomarkers, personalize therapies, and monitor neurological health and treatment response. The Dreem headband has the potential to scale the usage of a reduced montage PSG to the population at reasonable cost.

## AUTHOR CONTRIBUTIONS

Study concept and design: PJA, MEB, MG, MC, FS. Data acquisition: HJ, MH, PVB, MG, FS. Data analysis: VT, ABH, AG. Data interpretation: PJA, VT, FS. Writing the manuscript: PJA, VT, FS. Revising the manuscript: PJA, VT, MEB, FS.

## ACKNOWLEDGEMENTS

We would like to thank the Fatigue and Vigilance team including Drogou C., Erblang M., Dorey R., Quiquempoix M., Gomez-Merino D. and Rabat A. for their help in the study. We would like to thank Mignot E. for his help on the manuscript. We also would like to thank the Dreem team for their commitment to working on the Dreem headband over these years.

## CONFLICT OF INTEREST

PJA and MEB are employees of Dreem, Inc. and VT, ABH, AG, MH, and HJ of Dreem sas. FS was the principal investigator of the study. FS, MG and PVB declare that the research was conducted in the absence of any commercial or financial relationships that could be construed as a potential conflict of interest.

## FUNDING

This study was supported by Dreem sas.

